# Detection of vancomycin-resistant Enterococcus faecium and discrimination of vanA and vanB genotypes by VITEK MS Prime MALDI-TOF Mass Spectrometry and machine learning

**DOI:** 10.64898/2026.11.18.767493

**Authors:** María Sánchez-Cueto, Mark Gutierrez-Pareja, Emilia Cercenado, Patricia Muñoz, Mercedes Marín, Belén Rodríguez-Sánchez, David Rodriguez-Temporal

**Author notes:** Corresponding Authors: Belén Rodríguez-Sánchez and María Sánchez-Cueto Clinical Microbiology and Infectious Diseases Department, Hospital General Universitario Gregorio Marañón, Instituto de Investigación Sanitaria Gregorio Marañón (IISGM). Doctor Esquerdo, 46. 28007 Madrid, Spain.

## Abstract

Rapid identification of vancomycin-resistant Enterococcus faecium (VREfm) and discrimination of resistance mechanisms (VanA and VanB) are critical for appropriate antimicrobial therapy and infection control, particularly during outbreaks. Matrix-assisted laser desorption ionization–time of flight mass spectrometry (MALDI-TOF MS), combined with machine learning, has emerged as a promising tool for this purpose.

We evaluated whether MALDI-TOF mass spectra acquired on VITEK MS PRIME, combined with machine learning, could provide this information. Spectra from 388 from clinical and rectal surveillance samples (135 vancomycin-susceptible (VSEfm), 138 vanA and 115 vanB VREfm) were processed with Clover MSDAS, and four algorithms (partial least squares-discriminant analysis (PLS-DA), linear and polynomial support vector machines (SVM-L and SVM-P) and random forest (RF)) were trained to discriminate VSEfm from VREfm and VanA from VanB isolates. Model performance was evaluated using 10-fold cross-validation and using independent set of characterized E. faecium isolates from routine practice. Pairwise and triplet ensembles of these algorithms were also assessed.

In the test set, PLS-DA was the best individual classifier for VSEfm versus VREfm (balanced accuracy, 78.9%), and the ensemble of PLS-DA and linear SVM reached 80.7%. Validation within routine isolates confirmed the model’s robustness, achieving 100% correct classification. Discrimination of VanA from VanB was more accurate: 93.2% with RF and up to 97.1% with ensembles. Identical sequence types were found among susceptible, VanA and VanB isolates, indicating that classification was not driven by clonal lineage. MALDI-TOF MS combined with machine learning is a promising, low-cost tool for VREfm screening and vanA/vanB differentiation.

Importance: Vancomycin-resistant Enterococcus faecium is a leading cause of hospital-acquired infections. Laboratories usually need an additional day of susceptibility testing, or costly molecular assays, to detect resistance and determine whether it is mediated by vanA or vanB, information that guides treatment and infection control. MALDI-TOF mass spectrometry, already used in most clinical laboratories to identify bacteria within minutes. We have demonstrated that spectra, analysed using machine learning models on the VITEK MS PRIME platform, can also distinguish between vanA and vanB strains with high accuracy, without the need for additional reagents or manual handling time. Combining several algorithms improved reliability, and validation with genotyped isolates showed that results were not explained by the spread of clones.

This approach represents a rapid and cost-effective tool with potential clinical application in patient’s management and infection control. This approach could provide an early, low-cost alert at the time of identification, supporting earlier contact precautions and targeted therapy.

**Graphical abstract:** 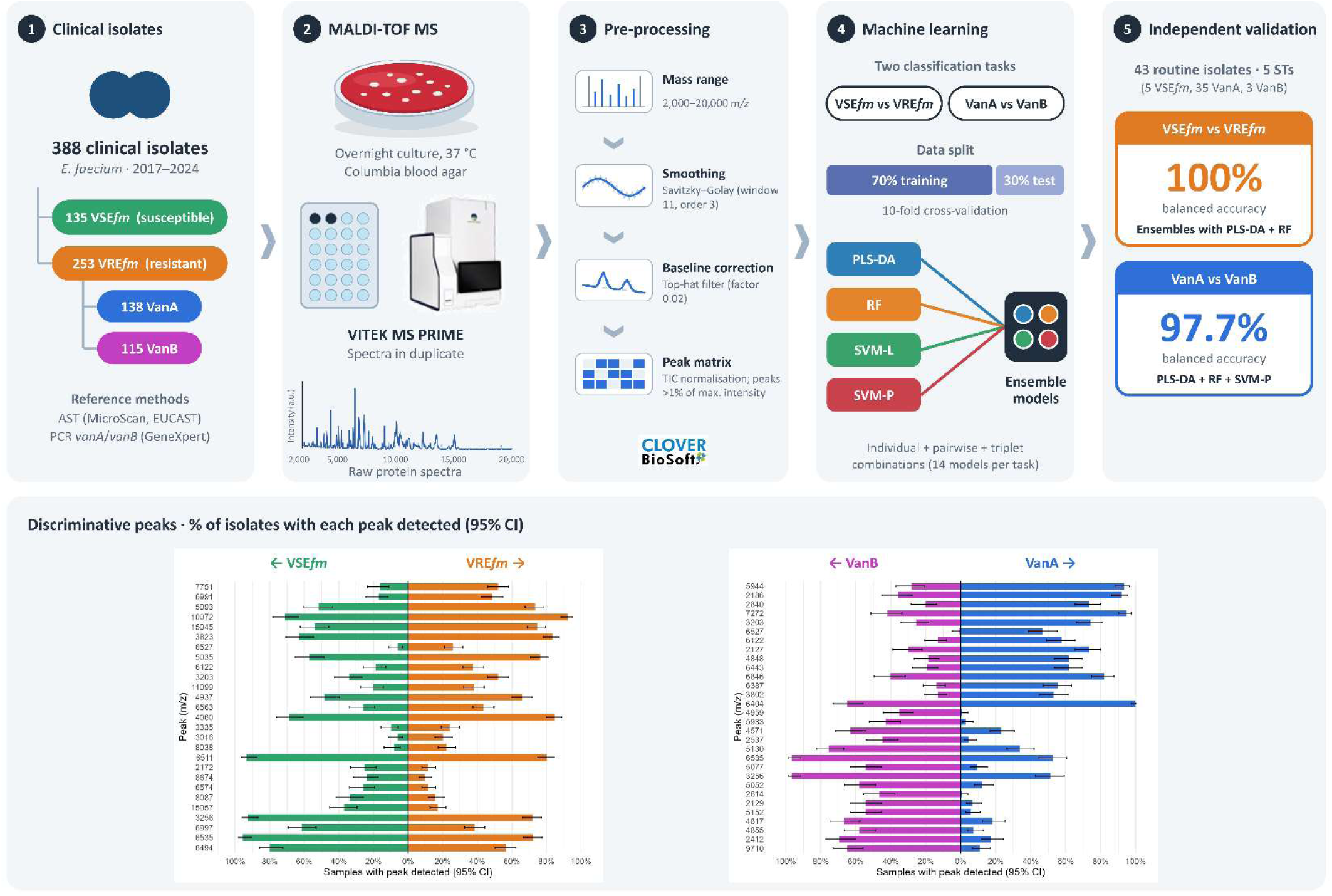

## Introduction

Enterococci are commensal bacteria that colonize the human gastrointestinal tract but can become opportunistic pathogens, particularly in critically ill and immunocompromised patients (1). The first clinical isolates of vancomycin-resistant enterococci (VRE) were reported in England in 1986. Since then, VRE has become widely disseminated, with reservoirs identified in both livestock and human populations across Europe (2). Transmission is primarily mediated through direct or indirect contact; furthermore, the remarkable genomic plasticity of these organisms facilitates their persistence in diverse and rigorous environments, particularly within clinical settings. Hospital-associated Enterococcus faecium strains are frequently multidrug-resistant and exhibit an enhanced capacity for adaptation and transmission within healthcare environments (3–5). As a result, VRE has emerged as a major global health concern, as recognized by the World Health Organization and the European Centre for Disease Prevention and Control (6, 7), and numerous hospital outbreaks of VREfm have been reported worldwide in recent years(8, 9)

Vancomycin susceptibility is routinely assessed in clinical microbiology laboratories using phenotypic methods, such as broth microdilution or gradient strip testing, which may take up to 24 hours. Alternatively, molecular techniques can identify vanA and vanB resistance determinants within 1 to 3 hours following initial culture growth (10). While these molecular assays significantly reduce turnaround time, their routine implementation is often constrained by higher operational costs (11).

Matrix-assisted laser desorption/ionization time-of-flight mass spectrometry (MALDI-TOF MS) is a rapid and accurate diagnostic tool that has become a standard in clinical microbiology laboratories, either replacing or complementing conventional phenotypic techniques for identifying most bacterial isolates (12, 13). In recent years, this technique has been proposed as a rapid and cost-effective method for detecting isolates hosting antimicrobial resistances. Specific biomarkers have been studied for bacteria from the ESKAPE group (14, 15), as well as specific resistance mechanisms, such as the detection of the mecA gene in methicillin-resistant Staphylococcus aureus (16, 17), β-lactamase production in Enterobacteriaceae, (18) and the cfiA gene in Bacteroides fragilis(19, 20). Recent studies on vancomycin resistance in Enterococcus have been published; however, the results exhibit considerable variability (11, 21–26). The application of MALDI-TOF MS in clinical microbiology laboratories holds promise as a rapid method for distinguishing vancomycin-resistant enterococci (VRE) from vancomycin-susceptible enterococci (VSE) by analyzing their unique protein mass spectra profiles.

The primary goal of this study was to develop a MALDI-TOF-based classification algorithm capable of differentiating vancomycin-susceptible E. faecium (VSEfm) from vancomycin-resistant E. faecium (VREfm), and to further classify VREfm isolates according to the presence of either the VanA or VanB resistance mechanism. As a secondary outcome, we aimed to validate these models using molecularly characterized E. faecium strains from routine practice. Additionally, an in silico protein peak analysis was performed to identify specific peaks useful for distinguishing resistant strains.

## Material and Methods

### 2.1. Bacterial Strains

A total of 388 E. faecium isolates recovered from clinical and rectal surveillance samples were included in the study (Table S1). They encompassed 135 VSEfm isolates and 253 VREfm. The resistant isolates were divided into 138 isolates harbouring the vanA and 115 the vanB gene. The isolates were obtained from clinical samples of patients admitted to the Hospital General Universitario Gregorio Marañón (HGUGM) in Madrid between 2017 and 2024, only one isolate per patient was included. Species-level identification was performed using MALDI VITEK® MS PRIME (bioMérieux, Marcy-l’Étoile, France). Following characterization, all strains were cryopreserved at -80°C for subsequent analysis. For this study, the samples were thawed and incubated overnight at 37°C on Columbia blood agar (bioMérieux).

### 2.2 Antibiotic Susceptibility Testing for Vancomycin

Antimicrobial susceptibility testing was performed as part of routine clinical diagnostics at the time of isolation, usingthe automated microdilution method Microscan® System (Beckman-Coulter, Brea, CA, USA) using PM33 panels following the manufacturer’s guidelines. Vancomycin breakpoints were established as indicated by the EUCAST (2025) v.15.0; the vancomycin breakpoint for enterococci remained unchanged throughout the study period. For isolates showing a vancomycin-resistant phenotype, the presence of resistance determinants was confirmed by commercial PCR GeneXpert® vanA/vanB cartridges (Cepheid, Sunnyvale, CA, USA).

### 2.3 Spectra acquisition by MALDI-TOF MS and data processing

Bacterial strains were analysed in duplicate using VITEK® MS PRIME (bioMérieux), according to the manufacturer’s instructions. A fresh colony was applied to a 48-well target plate and coated with 1 µL of α-cyano-4-hydroxycinnamic acid (CHCA) matrix solution. After drying, the target plate was inserted into the MALDI-TOF VITEK MS system. The generated MS fingerprint was automatically matched against the VITEK MS database v3.2. Escherichia coli ATCC 8739 was used as the quality control strain (27).

Protein spectra were processed with Clover MS Data Analysis software (Clover MSDAS; https://platform.clovermsdataanalysis.com, v1.15.12, Clover Biosoft, Granada, Spain). For classification analysis, peak matrices were generated within the 2,000– 20,000 m/z range. Spectral pre-processing included signal smoothing using a Savitzky– Golay filter (window length of 11 and polynomial order of 3), followed by baseline correction with the top-hat filter method (factor 0.02). Finally, replicates were grouped by sample name to obtain a single average spectrum per isolate.

### 2.4 Predictive models

Predictive models for the classification of VSEfm versus VREfm and to further differentiate VREfm isolates according to their VanA and VanB resistance mechanisms were developed using Clover MSDAS. Following spectral processing as described in Section 2.3, peak matrices were generated under the following conditions: intensities were then normalized by total ion current (TIC) normalization, and peaks were selected with intensities above 1.0% of the maximum peak intensity.

Four supervised Machine Learning algorithms were applied: Partial Least Squares-Discriminant Analysis (PLS-DA), Support Vector Machine linear (SVM-L) and polynomial (SVM-P), and Random Forest (RF). Internal validation of these models was performed using the K-fold cross-validation (K=10). For validation, total spectra were randomly split into a training set (70%) and a test set (30%).

For the VSEfm versus VREfm model, 271 isolates were used as training set (87 susceptible, 184 resistant) and 117 as a test set (48 susceptible and 69 resistant). For the VanA versus VanB model, 177 isolates were included in the training set (96 VanA and 81 VanB), and 76 were used as a test set (42 VanA and 34 VanB).

To assess whether predictive performance could be enhanced through model integration, an ensemble learning approach was implemented by combining the four individual classification models derived from a single peak matrix. The four algorithms (PLS-DA, SVM-L, SVM-P, and RF) were combined in all possible pairwise (n = 6) and triplet (n = 4) configurations, resulting in a total of 14 evaluated models and ensemble combinations (including the four individual models). For each combination, the definitive result was considered when all models agreed. Sensitivity, specificity, positive predictive value (PPV), and negative predictive value (NPV) were calculated using R software (version 4.4.2).

### 2.5 Validation with routine clinical isolates

The predictive models were further validated using 43 molecularly characterized E. faecium isolates obtained from routine laboratory practice, none of which had been included in the training or test sets (Table S2). This collection comprised two groups. The first included 25 isolates recovered between June and October 2024 from patients involved in a suspected nosocomial outbreak, all of which carried the vanA gene and were characterized by conventional multilocus sequence typing (MLST), based on PCR amplification and sequencing of seven housekeeping genes (28). Allelic profiles and sequence types (STs) were assigned using the PubMLST database (https://pubmlst.org/organisms/enterococcus-faecium). The second group consisted of 18 isolates (5 VSEfm, 10 VanA and 3 VanB VREfm) submitted to the laboratory for identification during the study period. They were characterized by whole-genome sequencing (WGS): libraries were prepared with the Nextera XT DNA Sample Preparation Kit (Illumina, Inc., San Diego, CA, USA) and processed on the Illumina MiSeq platform for MLST.

### 2.6 Protein peak analysis

An analysis was performed to identify potential biomarkers by leveraging group-specific protein m/z peaks as indicators for the precise classification of VREfm vs VSEfm and VanA vs VanB strains. First, all peaks were screened for tentative protein identification following previously described pipeline (29), which was implemented using Python (v3.13). Second, for differentiation of VREfm from VSEfm and VanA from VanB isolates, all peaks from the peak matrix created were analysed for comparison between groups following two approaches: 1) presence/absence of peaks was analysed by the proportion of samples showing each peak for each group and measuring the odds ratio and the Fisher test; 2) intensity of peaks was analysed comparing medians between each group for each peak and applying the Mann-Whitney test. P-values were adjusted for multiple comparisons using the Benjamini–Hochberg procedure to control the false discovery rate. Adjusted p-values < 0.05 were considered statistically significant. Statistical analyses were performed by R software.

## 3. Results

### 3.1. VSEfm vs VREfm classification models

Although RF algorithm showed the highest sensitivity (98.6%), this was associated with a markedly reduced specificity (35.4%), resulting in a lower balanced accuracy (Table 1). In contrast, PLS-DA and SVM-L algorithms maintained a higher balance between sensitivity (86.9% for both) and specificity (70.8% and 68.8%, respectively) (Table 1), with an Area Under the Curve (AUC) between 0.761-0.908 (Figure 1).

**Figure 1.**
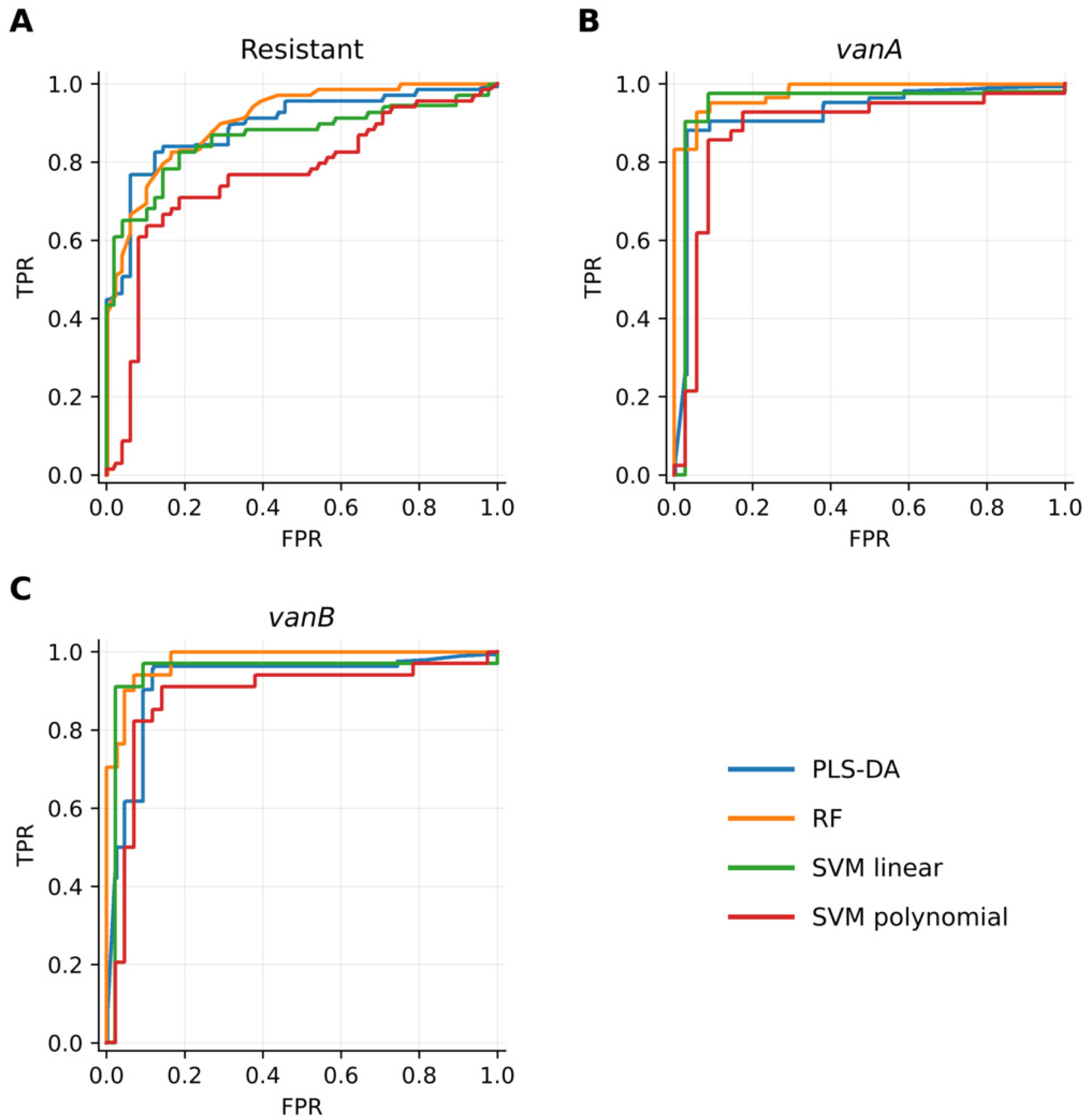
Receiving Operating Curve (ROC) for the positive detection of each category in both models (resistant model and vanA vs vanB model) for each algorithm applied. PLS-DA: Partial Least Squares-Discriminant Analysis; SVM: Support Vector Machine; RF: Random Forest.

**Table 1.** Results of the classification algorithms for the discrimination between vancomycin-susceptible and vancomycin-resistant Enterococcus faecium isolates.

| **Algorithm** | **Sensitivity** | **Specificity** | **PPV** | **NPV** | **Balanced accuracy** |
|---|---|---|---|---|---|
| PLS-DA | 87.0 | 70.8 | 81.1 | 79.1 | 78.9 |
| RF | 98.6 | 35.4 | 68.7 | 94.4 | 67.0 |
| SVML | 87.0 | 68.8 | 80.0 | 78.6 | 77.9 |
| SVMP | 94.2 | 20.8 | 63.1 | 71.4 | 57.5 |
| PLS-DA + RF | 98.4 | 56.0 | 84.5 | 93.3 | 77.2 |
| PLS-DA + SVML | 89.2 | 72.1 | 82.9 | 81.6 | 80.7 |
| PLS-DA + SVMP | 96.7 | 40.9 | 81.7 | 81.8 | 68.8 |
| RF + SVML | 98.4 | 53.6 | 82.2 | 93.8 | 76.0 |
| RF + SVMP | 98.5 | 11.1 | 73.0 | 75.0 | 54.8 |
| SVML + SVMP | 95.2 | 39.1 | 80.8 | 75.0 | 67.1 |
| PLS-DA + RF + SVML | 98.3 | 58.3 | 85.3 | 93.3 | 78.3 |
| PLS-DA + RF + SVMP | 98.3 | 23.1 | 85.3 | 75.0 | 60.7 |
| PLS-DA + SVML + SVMP | 96.6 | 45.0 | 83.8 | 81.8 | 70.8 |
| RF + SVML + SVMP | 98.3 | 20.0 | 83.1 | 75.0 | 59.2 |

PLS-DA: Partial Least Squares-Discriminant Analysis; SVML: Support Vector Machine linear; SVMP: Support Vector Machine polynomial; RF: Random Forest; PPV: Positive Predictive Value; NPV: Negative Predictive Value.

The pairwise ensemble models yielded a higher discrimination compared to individual models: the PLS-DA + SVM-L combination achieved the highest balanced accuracy (80.7%), with a sensitivity of 89.2% and a specificity of 72.1%. Other combinations generally increased sensitivity (up to 98.5%) but often at the expense of specificity, resulting in lower balanced accuracies.

Among the triplet ensembles, PLS-DA + RF + SVM-L yielded the highest balanced accuracy (78.3%), with high sensitivity (98.3%) and moderate specificity (58.3%). Overall, ensemble integration tended to enhance sensitivity but frequently reduced specificity, particularly in models including SVM-P, thereby limiting gains in balanced accuracy.

### 3.2. VanA vs VanB classification models

For the differentiation between VanA and VanB resistance mechanisms, RF achieved the highest balanced accuracy (93.2%), followed by SVM-L (92.6%) and PLS-DA (91.4%), whereas SVM-P showed comparatively lower performance (86.1%). RF demonstrated the highest sensitivity (95.2%), while PLS-DA and SVM-L achieved the highest specificity (97.1%; Table 2), with an AUC between 0.883-0.980 (Figure 1).

**Table 2.** Results of the classification algorithms for the discrimination between VanA and VanB Enterococcus faecium isolates.

| Algorithm | Sensitivity | Specificity | PPV | NPV | Balanced accuracy |
|---|---|---|---|---|---|
| PLS-DA | 85.7 | 97.1 | 97.3 | 84.6 | 91.4 |
| RF | 95.2 | 91.2 | 93.0 | 93.9 | 93.2 |
| SVML | 88.1 | 97.1 | 97.4 | 86.8 | 92.6 |
| SVMP | 92.9 | 79.4 | 84.8 | 90.0 | 86.1 |
| PLS-DA + RF | 97.2 | 96.9 | 97.2 | 96.9 | 97.1 |
| PLS-DA + SVML | 87.8 | 97.1 | 97.3 | 86.8 | 92.4 |
| PLS-DA + SVMP | 94.6 | 96.4 | 97.2 | 93.1 | 95.5 |
| RF + SVML | 97.3 | 96.9 | 97.3 | 96.9 | 97.1 |
| RF + SVMP | 97.4 | 90.0 | 92.7 | 96.4 | 93.7 |
| SVML + SVMP | 94.7 | 96.4 | 97.3 | 93.1 | 95.6 |
| PLS-DA + RF + SVML | 97.2 | 96.9 | 97.2 | 96.9 | 97.1 |
| PLS-DA + RF + SVMP | 97.1 | 96.4 | 97.1 | 96.4 | 96.8 |
| PLS-DA + SVML + SVMP | 94.6 | 96.4 | 97.2 | 93.1 | 95.5 |
| RF + SVML + SVMP | 97.2 | 96.4 | 97.2 | 96.4 | 96.8 |

PLS-DA: Partial Least Squares-Discriminant Analysis; SVML: Support Vector Machine linear; SVMP: Support Vector Machine polynomial; RF: Random Forest; PPV: Positive Predictive Value; NPV: Negative Predictive Value.

Pairwise ensemble combinations further improved performance. The PLS-DA + RF and SVM-L + RF models achieved the highest balanced accuracies (97.1%), maintaining both high sensitivity (97.2% and 97.3% respectively) and high specificity (96.9%). Other pairwise combinations also demonstrated balanced accuracies above 93%.

Among the triplet ensembles, SVM-L + PLS + RF achieved a balanced accuracy of 97.1%, with sensitivity and specificity of 97.2% and 96.9%, respectively. Overall, ensemble models consistently outperformed individual classifiers by achieving simultaneously high sensitivity and specificity, resulting in superior balanced accuracy.

### 3.3. Validation with routine clinical isolates

Among the 43 molecularly characterized E. faecium isolates from routine practice, 5 different sequence types (STs) were identified: 5 VSEfm (ST80) and 38 VREfm: 35 VanA (ST117=11, ST1421=14, ST80=6, ST78=2, ST17=2) and 3 VanB (ST117).

In the validation of the VSEfm versus VREfm classification model, the best performance was obtained with all ensemble combinations including PLS-DA and RF, which achieved a balanced accuracy of 100%. The results for the remaining models are summarised in Table 3 and Table S3.

**Table 3.** Results of the classification algorithms for the validation with routine clinical isolates between vancomycin-susceptible and vancomycin-resistant E. faecium isolates.

| Algorithm | Sensitivity | Specificity | PPV | NPV | Balanced accuracy |
|---|---|---|---|---|---|
| PLS-DA | 89.5 | 80.0 | 97.1 | 50.0 | 84.7 |
| RF | 94.7 | 80.0 | 97.3 | 66.7 | 87.4 |
| SVML | 92.1 | 80.0 | 97.2 | 57.1 | 86.1 |
| SVMP | 55.3 | 40.0 | 87.5 | 10.5 | 47.6 |
| PLS-DA + RF | 100 | 100 | 100 | 100 | 100 |
| PLS-DA + SVML | 97.0 | 100 | 100 | 75.0 | 98.5 |
| PLS-DA + SVMP | 87.0 | 66.7 | 95.2 | 40.0 | 76.8 |
| RF + SVML | 100 | 100 | 100 | 100 | 100 |
| RF + SVMP | 100 | 66.7 | 95.0 | 100 | 83.3 |
| SVML + SVMP | 95.0 | 100 | 100 | 50.0 | 97.5 |
| PLS-DA + RF + SVML | 100 | 100 | 100 | 100 | 100 |
| PLS-DA + RF + SVMP | 100 | 100 | 100 | 100 | 100 |
| PLS-DA + SVML + SVMP | 94.7 | 100 | 100 | 50.0 | 97.4 |
| RF + SVML + SVMP | 100 | 100 | 100 | 100 | 100 |

PLS-DA: Partial Least Squares-Discriminant Analysis; SVML: Support Vector Machine linear; SVMP: Support Vector Machine polynomial; RF: Random Forest; PPV: Positive Predictive Value; NPV: Negative Predictive Value.

For the VanA versus VanB differentiation model, the highest performance was achieved with the PLS-DA + RF + SVM-P combination, which yielded a balanced accuracy of 97.7%, with a sensitivity of 95.5% and a specificity of 100%. The validation results for the remaining models are summarised in Table 4 and Table S4.

**Table 4.**
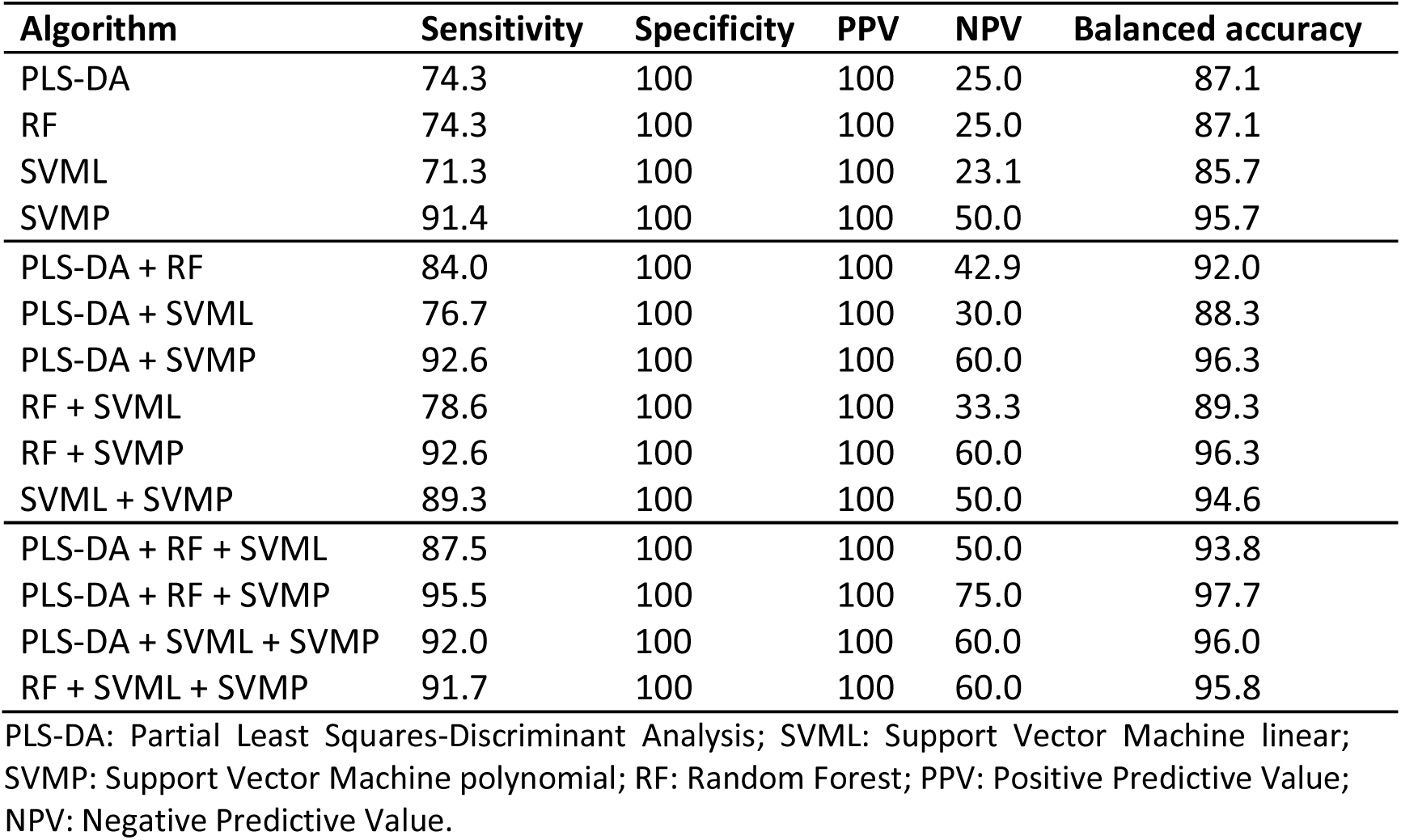
Results of the classification algorithms for the validation with routine clinical isolates between VanA and VanB E. faecium isolates.

| **Algorithm** | **Sensitivity** | **Specificity** | **PPV** | **NPV** | **Balanced accuracy** |
|---|---|---|---|---|---|
| PLS-DA | 74.3 | 100 | 100 | 25.0 | 87.1 |
| RF | 74.3 | 100 | 100 | 25.0 | 87.1 |
| SVML | 71.3 | 100 | 100 | 23.1 | 85.7 |
| SVMP | 91.4 | 100 | 100 | 50.0 | 95.7 |
| PLS-DA + RF | 84.0 | 100 | 100 | 42.9 | 92.0 |
| PLS-DA + SVML | 76.7 | 100 | 100 | 30.0 | 88.3 |
| PLS-DA + SVMP | 92.6 | 100 | 100 | 60.0 | 96.3 |
| RF + SVML | 78.6 | 100 | 100 | 33.3 | 89.3 |
| RF + SVMP | 92.6 | 100 | 100 | 60.0 | 96.3 |
| SVML + SVMP | 89.3 | 100 | 100 | 50.0 | 94.6 |
| PLS-DA + RF + SVML | 87.5 | 100 | 100 | 50.0 | 93.8 |
| PLS-DA + RF + SVMP | 95.5 | 100 | 100 | 75.0 | 97.7 |
| PLS-DA + SVML + SVMP | 92.0 | 100 | 100 | 60.0 | 96.0 |
| RF + SVML + SVMP | 91.7 | 100 | 100 | 60.0 | 95.8 |

PLS-DA: Partial Least Squares-Discriminant Analysis; SVML: Support Vector Machine linear; SVMP: Support Vector Machine polynomial; RF: Random Forest; PPV: Positive Predictive Value; NPV: Negative Predictive Value.

### 3.4. Protein peak analysis

All peaks observed in the spectra were analysed for search differences between susceptible and resistant isolates, and between VanA and VanB isolates. First, by the in silico analysis, we identified 68 peaks: 56 peaks assigned to ribosomal proteins and 12 to other proteins, as single and doubly charged, detected in most part of isolates (Table 5). For the differentiation of vancomycin resistance in our collection, 44 peaks were found to be strongly associated to these differences (Figure 2A): 27 showing statistical differences in appearance (Figure 2B) and 17 showing differences in intensity (Figure 2C). Among these peaks, some pairs were assumed to be the same protein as single and double charges, such as 5035-10072 m/z, 2548-5093 m/z, 3301-6601 m/z, 3256-6511 m/z and 2469-4937 m/z.

**Figure 2.**
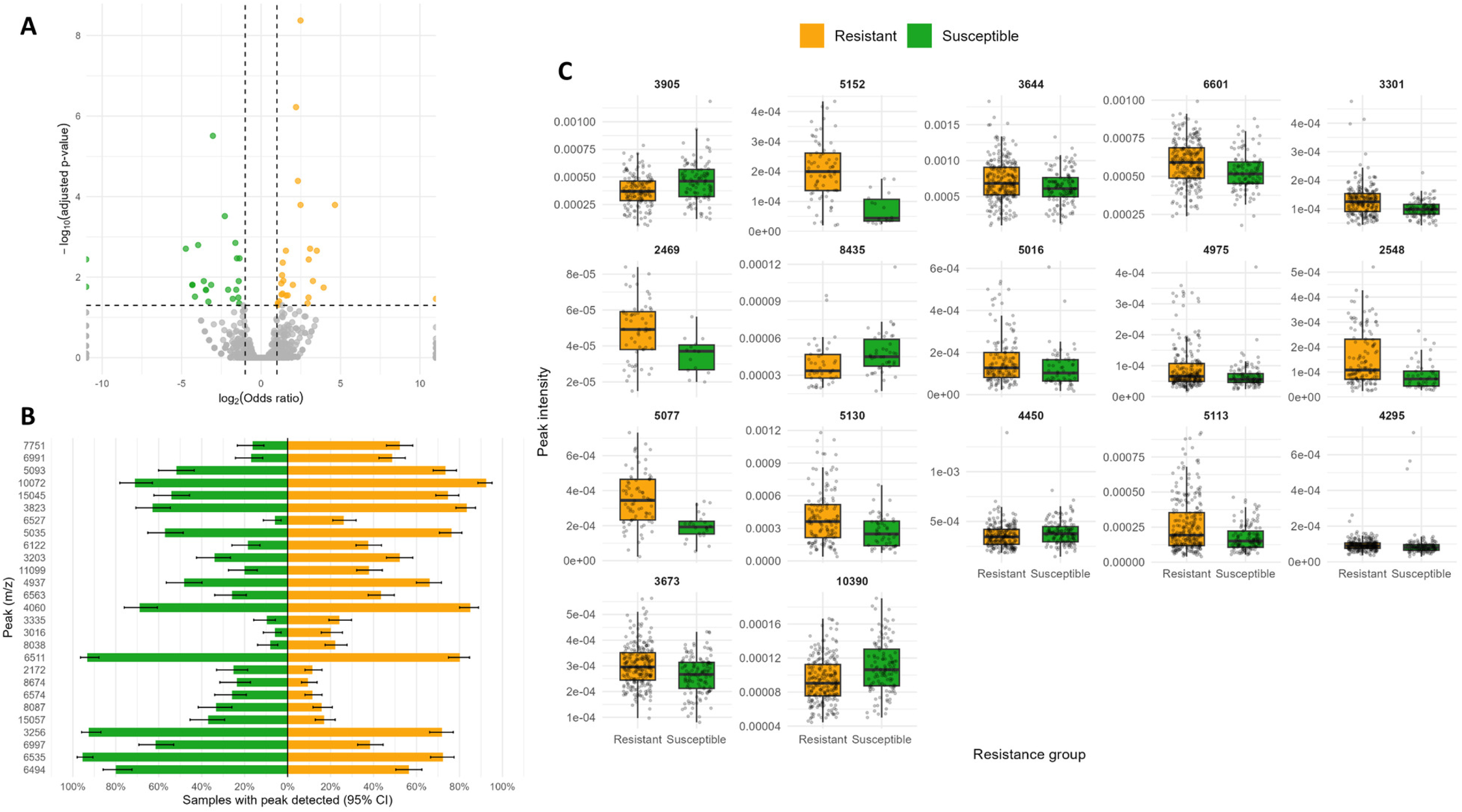
Discriminant peaks (m/z) between VREfm and VSEfm isolates. (A) Volcano plot of all peaks analysed. Vertical axe represents the statistical significance and horizontal axe represents the odds ratio for association with resistant (red) or susceptible (blue) categories. (B) Detection frequency (%) of each selected peak in vancomycin resistant (red) and susceptible (blue) isolates; error bars represent 95% confidence intervals. (C) Relative intensity of the peaks statistically significant in each group.

**Table 5.** Peaks identified by in silico analysis.

| **Peak (*m/z*)** | **Protein** | **Peak (*m/z*)** | **Protein** |
|---|---|---|---|
| 2,213 | RL36 [M+2H]<sup>2+</sup> | 6,512 | RL32-3 |
| 2,546 | Hiracin JM79 [M+2H]<sup>2+</sup> | 6,697 | Probable tautomerase EF_2859 |
| 2,678 | RL34 [M+2H]<sup>2+</sup> | 6,828 | RS21 |
| 3,165 | RL30-1 [M+2H]<sup>2+</sup> | 6,889 | RL28 |
| 3,172 | RL30 [M+2H]<sup>2+</sup> | 7,107 | RS9 [M+2H]<sup>2+</sup> |
| 3,256 | RL32-3 [M+2H]<sup>2+</sup> | 7,172 | Cold shock-like protein CspLA |
| 3,413 | RS21 [M+2H]<sup>2+</sup> | 7,289 | RS14-Z |
| 3,444 | RL28 [M+2H]<sup>2+</sup> | 7,328 | ATP-sintase subunit c |
| 3,645 | RS14-Z [M+2H]<sup>2+</sup> | 7,344 | RL29 |
| 3,672 | RL29 [M+2H]<sup>2+</sup> | 7,360 | RL11 [M+2H]<sup>2+</sup> |
| 3,842 | RL35 [M+2H]<sup>2+</sup> | 7,684 | RL35 |
| 4,059 | Translation initiation factor IF-1 [M+2H]<sup>2+</sup> | 8,119 | Translation initiation factor IF-1 |
| 4,428 | RL36 | 8,981 | RS18 |
| 4,493 | RS18 [M+2H]<sup>2+</sup> | 9,060 | RS20 |
| 4,530 | RS20 [M+2H]<sup>2+</sup> | 9,313 | Phosphocarrier protein Hpr |
| 4,656 | Phosphocarrier protein HPr [M+2H]<sup>2+</sup> | 9,552 | DNA-binding protein HU-1 |
| 4,777 | DNA-binding protein HU-1 [M+2H]<sup>2+</sup> | 9,951 | RL31-B |
| 4,975 | RL31-B [M+2H]<sup>2+</sup> | 10,004 | 10 kDa chaperonin |
| 5,094 | Hiracin JM79 | 10,120 | RS17 |
| 5,114 | RS16 [M+2H]<sup>2+</sup> | 10,224 | RS16 |
| 5,195 | RS19 [M+2H]<sup>2+</sup> | 10,391 | RS19 |
| 5,247 | RS15 [M+2H]<sup>2+</sup> | 10,492 | RS15 |
| 5,354 | RL34 | 10,938 | RL24 |
| 5,469 | RL24 [M+2H]<sup>2+</sup> | 11,041 | RL23 |
| 5,503 | RL33-2 | 11,101 | RL21 |
| 5,520 | RL23 [M+2H]<sup>2+</sup> | 11,563 | RS10 |
| 5,550 | RL21 [M+2H]<sup>2+</sup> | 12,319 | RL22 |
| 5,782 | RS10 [M+2H]<sup>2+</sup> | 12,324 | RL7/RL12-1 |
| 6,010 | RL33-3 | 12,338 | RL7/RL12 |
| 6,159 | RL22 [M+2H]<sup>2+</sup> | 12,980 | RL18 |
| 6,168 | RL7/RL12 [M+2H]<sup>2+</sup> | 13,403 | RS13 |
| 6,327 | RL30-1 | 13,490 | RL20 |
| 6,341 | RL30 | 14,214 | RS9 |
| 6,493 | RL18 [M+2H]<sup>2+</sup> | 14,717 | RL11 |

RL: ribosomal large; RS: ribosomal small.

Regarding the differentiation of resistance mechanisms (VanA vs VanB), 50 peaks were found to be strongly associated to these differences (Figure 3A): 30 showing statistical differences in appearance (Figure 3B) and 20 showing differences in intensity (Figure 3C). Among these peaks, some pair were also assumed to be the same protein as single and double charge: 3203-6406 m/z, 2537-5074 m/z, 2211-4429 m/z, 3644-7288 m/z and 4855-9710 m/z. Interestingly, the peak 6527 m/z showed to be specific for VanA (present in 46% of VanA isolates and 0.8% in VanB). On the other hand, the peak 4959 m/z (present in 35% of VanB isolates and 0.7% in VanA) and the peak 2613 m/z (present in 46% of VanB isolates and 0.7% in VanA) showed to be more specific for VanB (Figure 3B).

**Figure 3.**
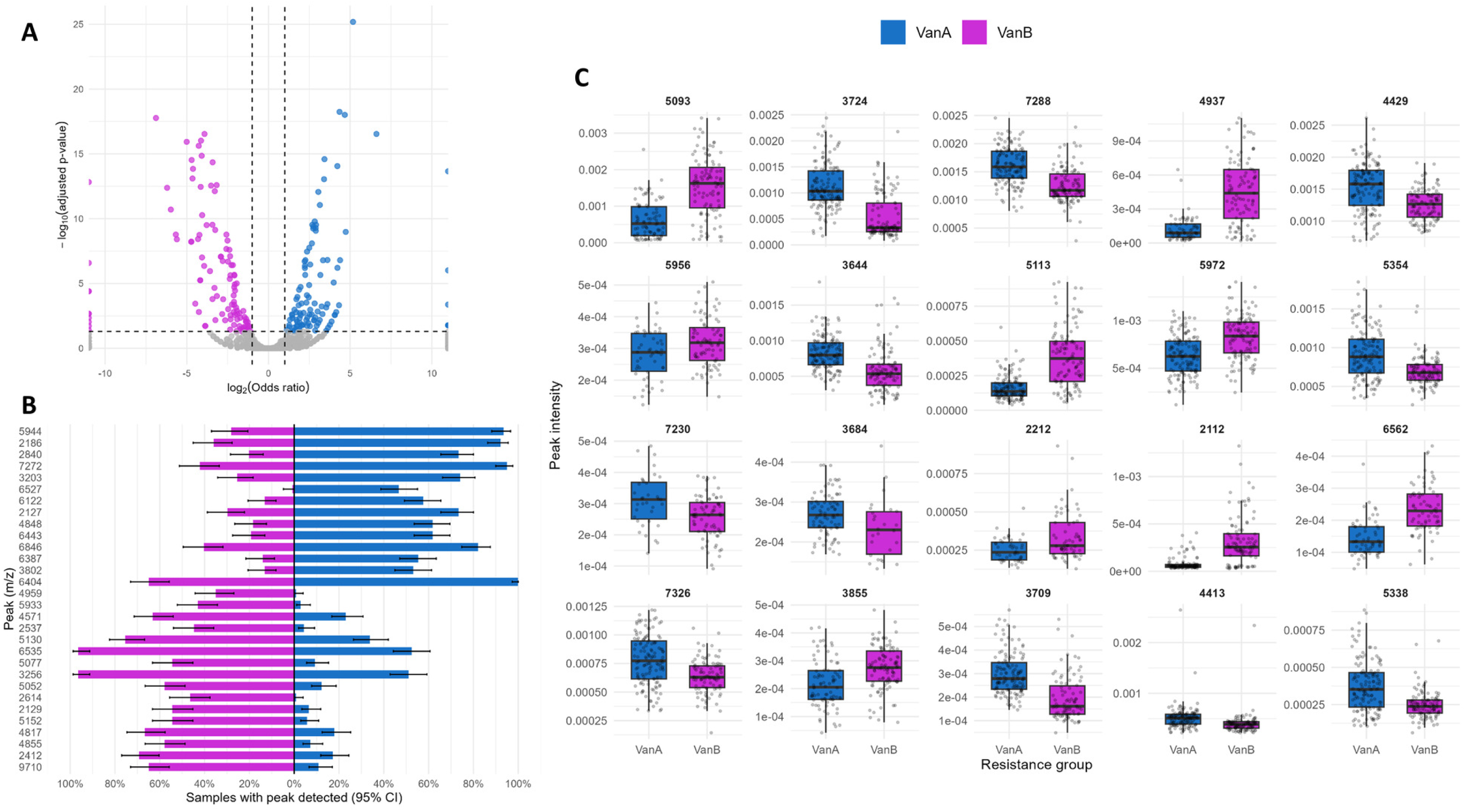
Discriminant peaks (m/z) between VanA and VanB VREfm isolates. (A) Volcano plot of all peaks analysed. Vertical axe represents the statistical significance and horizontal axe represents the odds ratio for association with VanA (red) or VanB (blue) categories. (B) Detection frequency (%) of each selected peak in VanA (red) and VanB (blue) isolates; error bars represent 95% confidence intervals. (C) Relative intensity of the peaks statistically significant in each group.

## Discussion

In this study, we have developed a classification model based on MALDI-TOF MS spectra processed with Clover MSDAS for the identification of VREfm and further classification of these isolates according to the underlying resistance gene, vanA or vanB. Individual classifiers already yielded promising results for the discrimination between susceptible and resistant isolates and ensemble models further improved performance, reaching a balanced accuracy of 100% in the validation set with combinations including PLS-DA and RF. For VanA/VanB differentiation, the PLS-DA + RF + SVM-P ensemble achieved a balanced accuracy of 97.7%.

For the discrimination between vancomycin-susceptible and vancomycin-resistant E. faecium, the PLS-DA algorithm achieved a correct classification rate of 78.9%, which is consistent with previous studies evaluating MALDI-TOF MS based machine-learning approaches for the detection of vancomycin resistance. Candela et al. reported accuracies ranging from 77.5% to 80.9% depending on the algorithm employed (11), and Gao et al., using the using the previous-generation VITEK MS system, obtained comparable accuracies of 78.07% to 80.50% (26). Wang et al. demonstrated the feasibility of predicting vancomycin resistance directly from MALDI-TOF MS spectra using large spectral datasets, achieving AUC values above 0.84 (24). Similarly, Griffin et al. demonstrated that vanB-positive E. faecium could be reliably differentiated from vanB-negative isolates using MALDI-TOF MS, their genetic algorithm achieved a recognition capability of 95.12% and a cross-validation accuracy of 88.24% (30). Likewise, Nakano et al. reported a discrimination rate of 92.4% for VanA-VRE isolates versus VSE isolates using a genetic algorithm-based model (22). Together, these studies support the growing utility of MALDI-TOF MS combined with machine-learning algorithms as a rapid strategy for resistance prediction in E. faecium.

When performance metrics beyond overall accuracy were considered, PLS-DA and SVM-L demonstrated the most balanced behaviour in terms of sensitivity and specificity, yielding the highest balanced accuracy values among individual classifiers. Although RF achieved very high sensitivity, this was occasionally associated with reduced specificity, potentially limiting its clinical reliability if used alone. To overcome these limitations, we implemented ensemble learning strategies, an approach that has previously been shown to improve the performance of MALDI-TOF MS-based classification models by combining the strengths of individual algorithms (31). Combinations integrating PLS-DA, SVM-L, and RF improved the equilibrium between sensitivity and specificity, resulting in higher balanced accuracy than individual models alone and suggesting that model integration enhances robustness for detecting vancomycin resistance. Most notably, validation with routine clinical yielded a correct classification rate of 100%, confirming strong generalisability in routine diagnostic conditions. These findings are comparable to those reported by Griffin et al., whose prospective validation integrated into routine laboratory workflows achieved a sensitivity of 96.7% and a specificity of 98.1% (30) , reinforcing the translational potential of MALDI-TOF MS–driven predictive models for rapid detection of vancomycin resistance.

The discrimination between vanA and vanB genotypes yielded consistently high performance across all algorithms with balanced accuracies ranging from 86.1% to 93.2% for individual classifiers and exceeding 95.0% for most ensemble combinations. These results indicate that MALDI-TOF MS spectral differences between VanA- and VanB-type isolates are more pronounced than those separating susceptible and resistant strains. One possible explanation for this finding may lie in the distinct genomic organization of these resistance determinants. The vanA genotype is commonly associated with the Tn1546 transposon, which generally exhibits a relatively conserved genetic structure, whereas vanB-associated elements are characterized by greater structural heterogeneity and more variable expression patterns. These differences could contribute to the generation of more reproducible MALDI-TOF MS spectral signatures among vanA isolates, thereby facilitating discrimination between both resistance genotypes (32).

A major limitation of the initial phase of the study was the inability to determine whether the discriminatory performance of the models was driven by resistance associated features or by clonal relatedness among isolates, since all strains were collected from a single healthcare centre. To address this, we conducted our second objective: the validation of the models using molecularly characterized, routine clinical isolates. During these validation, MLST analysis demonstrated that isolates displaying different resistance profiles were not clonally related, as identical STs were identified among both susceptible isolates and those resistant to VanA and VanB. Furthermore, the ensemble combinations correctly classified 100% of isolates in the VSEfm vs VREfm analysis and 97.7% in the VanA vs VanB differentiation model, demonstrating a discrimination based on the resistance mechanisms present, not on the clones.

Several peaks showed strong association with VSEfm or VREfm isolates, most notably those at 7751, 6991 and 6527 m/z, none of which has been previously reported in this context. These peaks were detected more frequently among resistant than susceptible isolates, although their presence in only a fraction of resistant isolates suggests that they are more useful in combination than as individual markers. Regarding peaks previously described in the literature, the peak at 5093 m/z was detected in 74% of resistant and 52% of susceptible isolates in our cohort. Griffin et al. had already reported a peak at 5094 m/z as a significant marker for the identification of VRE (30), and Candela et al. detected a peak at 5095 m/z in 82 of their 86 VREfm isolates (11). Brackmann et al. subsequently sequenced the 5095 m/z protein and identified it as hiracin, a secretory protein encoded by the hirJM79 gene whose role in vancomycin resistance remains unknown (33).

Regarding the differentiation between resistance genotypes, several peaks showed significant differences between VanA and VanB isolates. The peaks at 4959 and 2613 m/z were detected almost exclusively in VanB isolates, whereas the peak at 6527 m/z, was detected only in VanA isolates.

Despite these promising findings, several limitations should be acknowledged. First, all isolates included in the initial analysis were obtained from a single healthcare centre, and the predictive models were not validated simultaneously using multicentre collections. Consequently, the generalizability of the models across different epidemiological settings remains to be fully established. Second, although validation was performed with molecularly characterized routine clinical isolates, strain diversity was limited, particularly due to the predominance of vanA isolates in the study collection. In addition, isolates simultaneously harbouring both vanA and vanB resistance determinants were not included in this study, and therefore the ability of the models to classify strains carrying both mechanisms could not be evaluated. Nevertheless, this validation was sufficient to demonstrate that the discriminatory performance of the models was not explained by clonal relatedness among isolates.

On the other hand, this study has notable strengths. To our knowledge, it is the first to apply ensemble models combining heterogeneous classifiers (PLS-DA, SVM and RF) for VREfm detection. Because each algorithm makes different errors, their combination compensated for individual weaknesses. As a result, the ensembles achieved simultaneously high sensitivity and specificity, a key requirement for clinical use, without any additional laboratory work or cost. Finally, these machine-learning models require continuous refinement through the incorporation of a larger number of genetically diverse strains from multiple institutions in order to further evaluate their robustness, reproducibility, and potential applicability in routine clinical microbiology laboratories.

In conclusion, MALDI-TOF MS combined with machine-learning algorithms constitutes a reliable and rapid approach for the detection of vancomycin resistance in E. faecium and for the differentiation of its underlying genetic determinants (vanA and vanB genes). While classification of susceptible and resistant isolates showed moderate performance, VanA and VanB isolates were distinguished with higher accuracy, suggesting that spectral differences associated with each resistance genotype are more consistent than those separating resistant from susceptible isolates The implementation of ensemble learning strategies further improved model robustness by optimizing the balance between sensitivity and specificity. Importantly, external validation and MLST analysis confirmed that the observed discriminatory capacity was not driven by clonal relatedness, but may be attributable to characteristics associated with resistance. Altogether, these findings support the potential clinical utility of MALDI-TOF MS based machine-learning approaches as rapid and cost-effective screening tools for the early detection of vancomycin-resistant E. faecium. Because MALDI-TOF MS is already part of the routine workflow in most clinical microbiology laboratories, this approach could be implemented without additional reagents or hands-on time, as the same spectra acquired for species identification would be used for resistance prediction. Their implementation in routine clinical microbiology workflows could enable prompt identification of potentially resistant isolates, thereby supporting early therapeutic decision-making, antimicrobial stewardship, and infection control interventions.

## Data availability statement

Data used in this study can be requested from the corresponding author upon reasonable request.

